# Whole-tree heartwood inducing technique for *Dalbergia odorifera*

**DOI:** 10.1101/503987

**Authors:** Hui Meng, DeLi Chen, Xiangsheng Zhao, Zheng Zhang, Yangyang Liu, Zhihui Gao, Peiwei Liu, Jian Feng, Yun Yang, JianHe Wei

## Abstract

The dried heartwood of *Dalbergia odorifera* is a very important traditional Chinese medicine. In nature, *D. odorifera* heartwood forms slowly over several decades, and excessive harvesting has caused large-scale reductions of *its* wild populations. To date, there have been no studies on artificial methods to induce heartwood formation in trees that have not already begun producing heartwood. We have published a patent for a whole-tree heartwood inducing technique for *D. odorifera* (the “Dodor-Wit” method); here, we describe and analyze its efficacy. Trees in four plantations in China were treated using this method. After two years, in 2016 and 2017, they were harvested and analyzed. Average heartwood yield per tree was 3,161.95 g. Heartwood induction rate (dry heartwood weight to total dry trunk weight after peeling) was 21.60%. Average volatile oil content per tree was 1.42%. Alcohol-soluble extract of the induced heartwood was 15.77%, exceeding the Chinese Pharmacopoeia standard. This is the first report on the Dodor-Wit method and heartwood induction in *D. odorifera* trees aged 5–8 years. The heartwood induced by Dodor-Wit satisfies the requirements of the Chinese Pharmacopoeia. The Dodor-Wit method has an important role in solving the extreme scarcity of Chinese medicinal herbs.

## Introduction

There are 29 species of rosewood, of which 17 belong to the genus *Dalbergia.Dalbergia* spp. are mainly distributed in tropicaland subtropical regions. The most valuable rosewood species is *Dalbergia odorifera*T. Chen, which is endemic to Hainan Province, China[1]. The heartwood (resinous wood) of *D. odorifera* is used to produce high-value furniture, fragrances, and traditional Chinese medicine, for which it is referred to as “Jiang Xiang” [2,3]. Itsheartwood is also widely used in drug therapy in Japan, Singapore, and South Korea. Jiang Xiang, which is listed in the Chinese Pharmacopoeia, has medicinal effects for treating conditions such asblood disorders, ischemia, inflammation, necrosis, and rheumatic pain[4]. Jiang Xiang is used in Chinese herbal prescriptions such as *Qi-Shen-Yi-Qi* decoction[5], *Ten Fragrant Ingredients Pain-Re* [6], and *Guan-Xin-Dan-shen* pills[7]. There are 112 traditional Chinese medicine formulations containing Jiang Xiang, of which 21 are mentioned in the Chinese Pharmacopoeia. The annual demand for raw *D. odorifera* material is over 300 tonnes, and the annual production value exceeds 5 billion yuan. Because of its economic value and rarity, indiscriminate cutting of trees and overharvesting to obtain heartwood has depleted wild tree populations. *Dalbergia odorifera* was listed in the IUCN Red List of Threatened Species as an endangered species[8], and in 2017 was included in Appendix II of the Convention on International Trade in Endangered Species of Wild Fauna and Flora. Only a few wild *D. odorifera* trees remain, and these are found in nature reserves in Hainan province of China. *Dalbergia odorifera* for the Chinese medicine market now comes from heartwood that is retrieved from old furniture, handicrafts, and other wooden products that were originally made from wood collected in the wild; there is also existing stock held by Chinese medicine factories. The supply of *D. odorifera*heartwoodfrom wild sources is far less than the market demand.

Efforts have been made to preserve natural *D. odorifera* populations and to increase heartwood supply, including developing the cultivation of *D. odorifera* [9]. *Dalbergia odorifera* planting has taken place since the 2000s in the provinces of Hainan, Guangdong[10], Guangxi[11], Yunnan[12], Fujian[13], Jiangxi[14], Zhejiang[15], and Sichuan[16]. Several million *D. odorifera* trees have been cultivated in the last decade. *Dalbergia odorifera* trees grow slowly; their heartwood formation occurs unpredictably, and by inches. Heartwood typically begins to form 7–8 years after seedlingsare planted and can be used for wood processing at least 30 years after planting [17]. Therefore, trees that were planted in the 2000s are not yet able to provide heartwood for the wood and medicine markets.

In the wild, heartwood forms mostly in the center of the trunk and root. During artificial heartwood induction, the accumulation of heartwood substances gradually accumulates with time, and the color of the wood darkens. External factors such as environmental stress [18–20], mechanical injury[21–22], phytohormones [22–26], and insect attack or microbial invasion[27–29] cause the xylem to form heartwood around the wounded or rotting parts of the trunk or root. To date, there have been no reports of artificial heartwood induction in trees.

We have successfully induced agarwood formation in *Aquilaria sinensis* using the Agar-Wit technology[30]. This has been applied to 265,000 trees in China and other southeast Asian countries. Agar-Wit has become the most widely applied agarwood induction method. Agarwood has been widely used to produce medicine and incense, and its production conforms to the standard of the Chinese Pharmacopoeia. We have now developed a method to induce medicinal-quality heartwood in*D. odorifera*; this method is called “the whole-tree heartwood-inducing technique from *D. odorifera*” (hereafter “Dodor-Wit”)[22], and we have patented it in China.

Our objective in this study was to assess the induction capability of this method, and to evaluate the quality of the heartwood according to Chinese Pharmacopoeia standards [4]. To do this, we applied the Dodor-Wit method to trees at locations in China: Luoniushan plantation (LNS) in Haikou; Lelai plantation (LL) in Wanning; Jianfeng plantation (JF) in Ledong; and Banqiao plantation (BQ) in Dongfang.

## Results and discussion

### Induction process and formation of heartwood

According to the infusion model, dye solution was used to track the distribution of the induction solution, and the process and distribution of heartwood formation after the induction were indirectly demonstrated. We cut cross sections of the trees every 30 cmalong the trunk. As the plants transpired, the induction solution entered the tree at the infusion hole and was transmitted up the trunk to a height of 2–3 m depending on the tree. The dyed area gradually increased away from the infusion hole, and was clearly stained at 0.3 m to 3.3 m (Fig 1 (panels 2 to 11), and then decreased gradually. The induced heartwood occurred mainly at a height of 0.9 m to 2.1 m.

**Fig 1.**
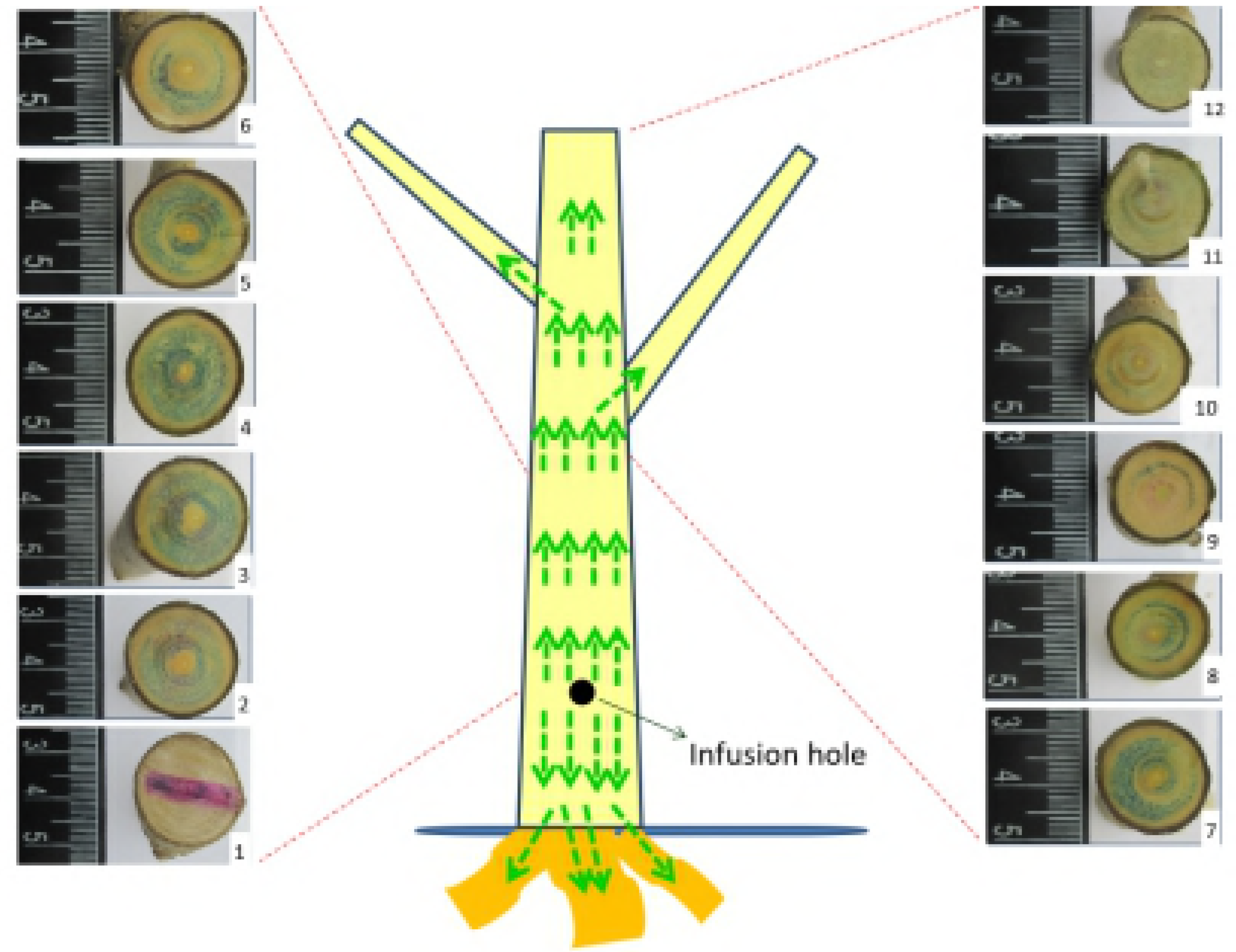
Infusion model of the Dodor-Wit method to induce heartwood formation in *Dalbergia odorifera*. **Panels 1–12:** Cross-sections that were taken every 30 cm above the infusion hole. The green dye indicates the distribution of the inducer.

A brown area and a thin resinous layer appeared inside the trunk within the first week after treatment. Puce-colored wood and a thick resinous layer were observed throughout the trunkafter 12 months. In the present study, we found that black resinous wood formed within24 months of induction treatment (Figs 2 and 3), and this wood reached the quality standards of the Chinese Pharmacopoeia. Therefore, the method successfully induced resinous wood production throughout the entire tree trunk.

**Fig 2.**
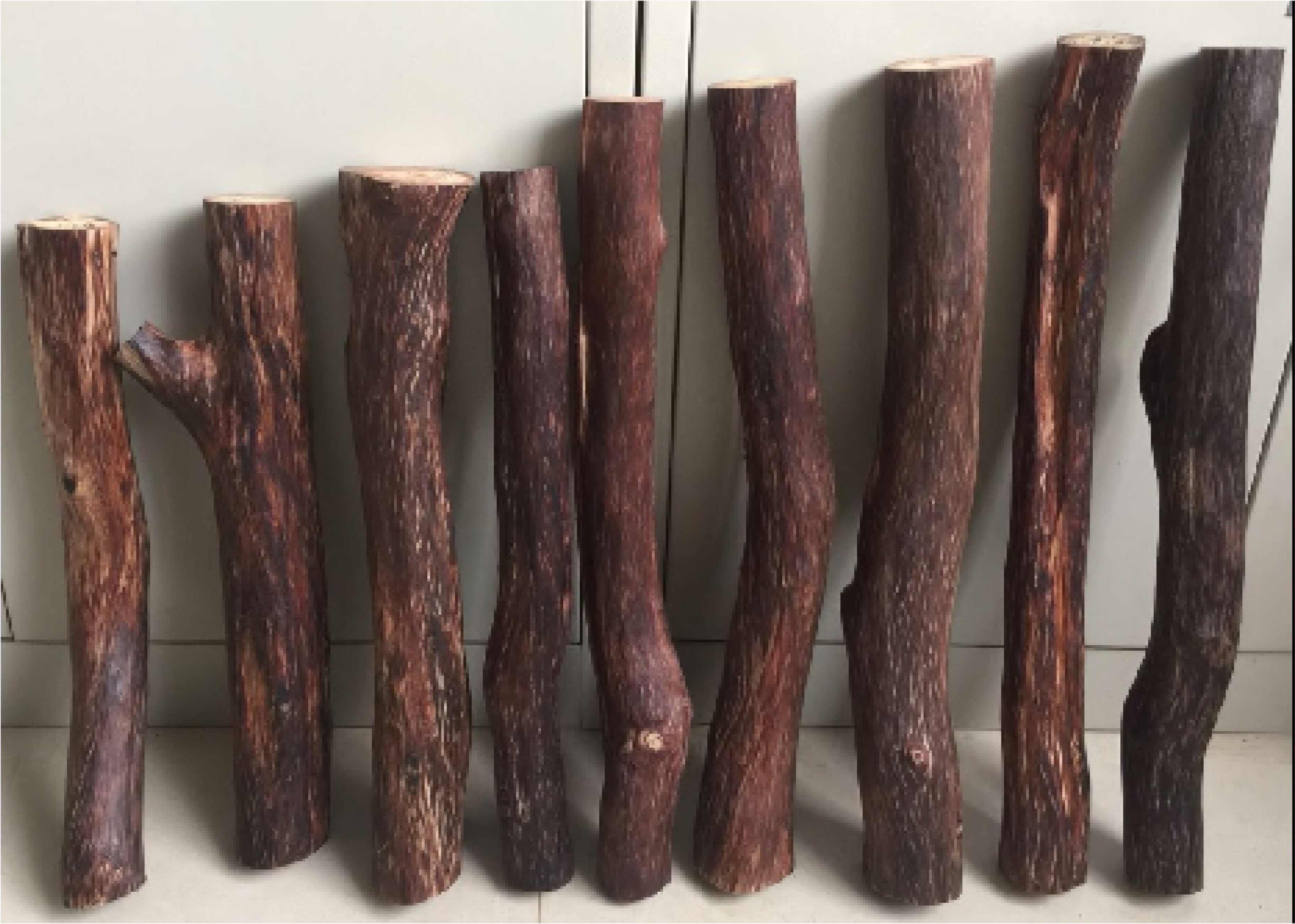
The heartwood produced by the Dodor-Wit method from *Dalbergia odorifera* tree trunks. The bark and sapwood have been peeled off.

**Fig 3.**
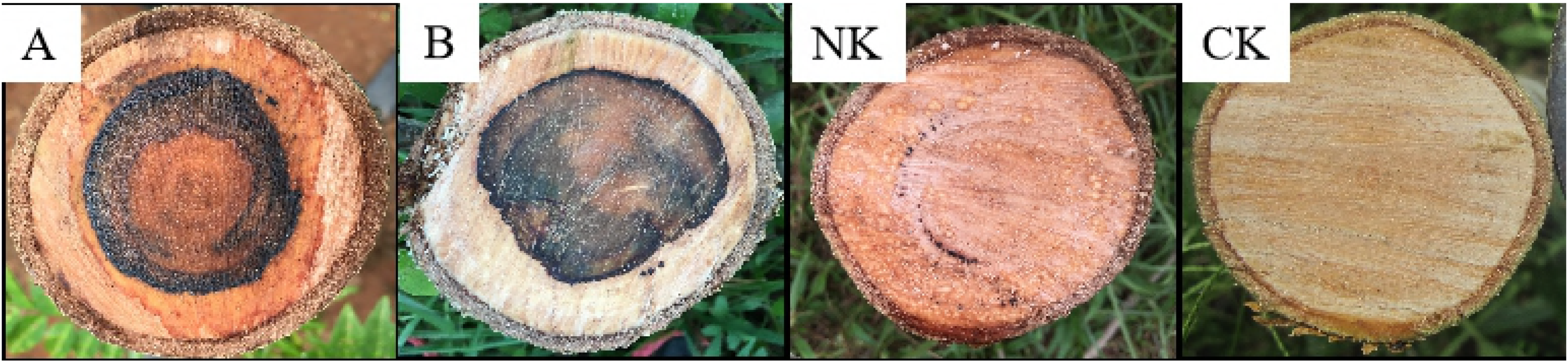
Cross-sections of *Dalbergia odorifera* trunks treated with the Dodor-Wit method to induce heartwood formation. A, B: wood treated with the Dodor-Wit method. NK: pure water negative control. CK: blank control of healthy wood. The trees were cultivated at Luoniushan plantation in Haikou, in the Hainan province of China.

### Tree mortality

The average annual mortality rate for *D. odorifera* trees treated using the Dodor-Wit method in the four plantations was only 3.78%(Table 1). We found that the causes of death after induction might be related to factors such as weather, topography, individual differences, and differences in how the trees are farmed at the different plantations.

**Table 1.** *Dalbergia odorifera* tree mortality in the four plantations (sites)after application of the Dodor-Wit method to induce heartwood formation.

| Site | Mortality in first year | Mortality in second year | Number of trees treated | Mortality rate* |
| --- | --- | --- | --- | --- |
| LNS | 1 | 4 | 70 | 7.14% |
| LL | 2 | 3 | 100 | 5.00% |
| JF | 0 | 0 | 100 | 0.00% |
| BQ | 4 | 0 | 100 | 4.00% |

\* The mortality rate is deduced by dividing the total number of plants by the number of deaths. LNS, LL, JF, and BQ refer to Luoniushan plantation, Haikou; Lelai plantation, Wanning; Jianfeng plantation, Ledong; and Banqiao plantation, Dongfang.

### Yield of induced heartwood in the trunk

The heartwood yield of each tree two years after induction ranged from 1,142.8 g to 4,707.3 g (Table 2). No resin formed in the natural untreated samples of these plantations. The heartwood induction rate (dry heartwood weight to total dry trunk weight after peeling) was 21.60%.

**Table 2.** Yield of induced heartwood two years after application of the Dodor-Wit method induce heartwood formation in *Dalbergia odorifera*. The trees were cultivated in China. The results are presented as means ± SD.

| Site | Average yield per tree*(g) | Number of samples | Total weight of samples(kg) |
| --- | --- | --- | --- |
| **LNS** | 3,297.4 ± 327.9 | 27 | 85,732.5 |
| **LL** | 1,142.8 ± 106.4 | 26 | 30,855.0 |
| **JF** | 4,707.3 ± 359.1 | 33 | 155,340.2 |

\* The average values are based on the number of samples in each plantation. Total weight of samples is the total weight of induced heartwood for each plantation. Each sample was tested three times. LNS, LL, and JF refer to Luoniushan plantation, Haikou; Lelai plantation, Wanning; and Jianfeng plantation, Ledong.

## Quality of induced heartwood

### Microscopic characteristics

Examination of the wood sections under the microscope revealed that the basic structure of natural heartwood and induced trees was consistent, and that they differed only in cell size. Further, it revealed that there was no difference in xylem structure before and after induction. The xylem had large and clear bordered pit vessels, which contained a reddish-brown or yellow-brown substance in the lumen after induction. Ray cells were in a single column, and rarely in two columns. Patches of pigment were reddish-brown or yellowish. The cell structure and heartwood characteristics of the trees after induction conformed to the description provided in the Chinese Pharmacopoeia.

The basic structure of the wood powder of natural heartwood and induced samples was similar. In these two panels, the lowercase letters (a, b, and c), and arrowsrefered to resins, fibers and ducts, respectively. The color of the two sampleswas also similar. Some tissue size differences could be caused by the age of the tree (Fig 4).

**Fig 4.**
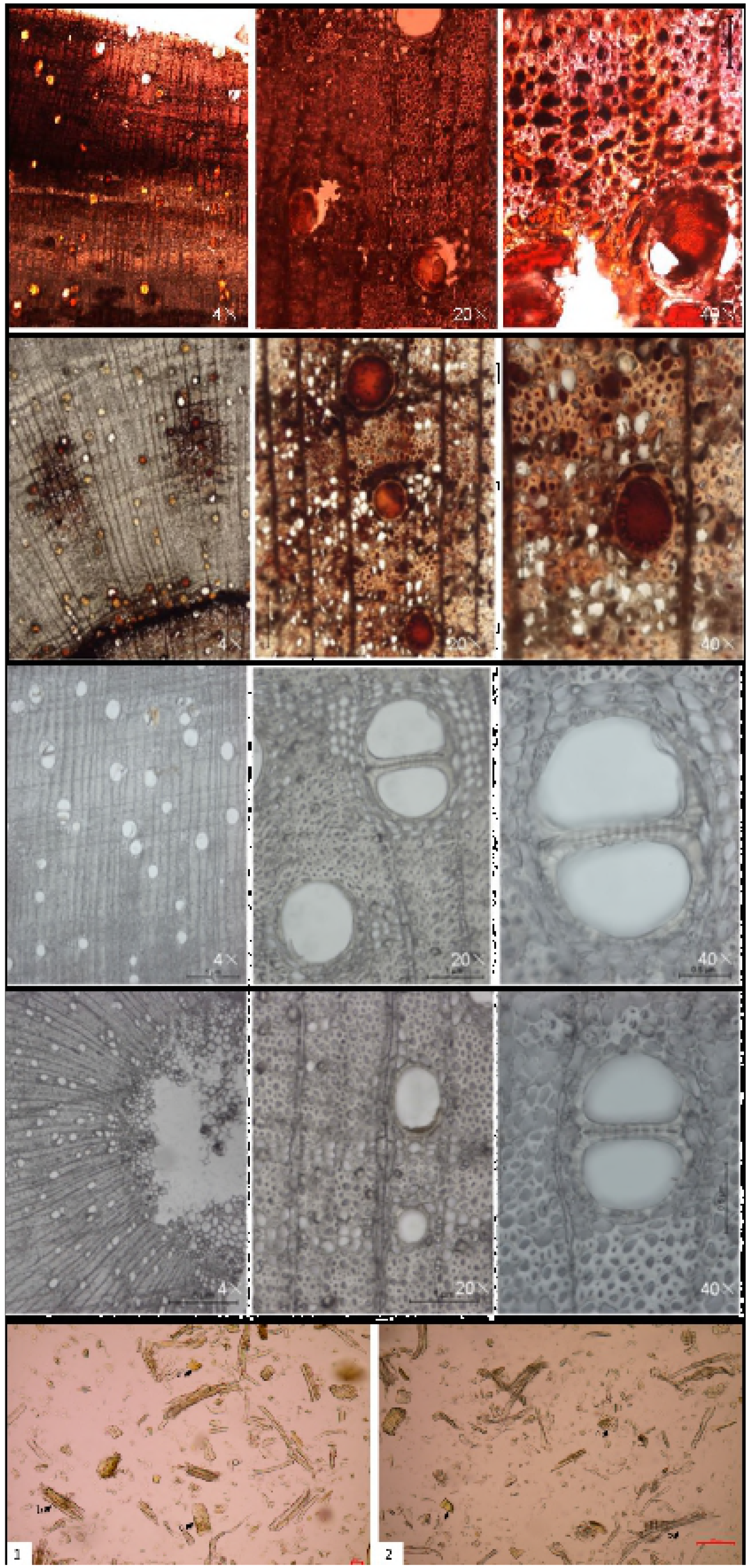
Microscopic characteristics of *Dalbergia odorifera* wood after application of the Dodor-Wit method. (A) natural heartwood (HW treatment); (B) Three-year-old tree, 6 months after it was treated usingthe Dodor-Wit method to induce heartwood formation; (C) Sapwood of natural heartwood (HW treatment); (D) Normal 3-year-old tree without treatment (CK treatment); (E) Wood powder (E1: natural heartwood; E2: induced heartwood).

### Identification of heartwood by TLC

The TLC fingerprint profiles of induced wood treated by the Dodor-Wit method samples 1-13 were compared with nerolidol (14) and the wild heartwood control (1,15) (Fig 5). The results displayed that there are five spots in common (*R*_*f*_ = 0.88, 057, 0.50, 0.41, 0.31) which appeared as blue bands, and threespots in common (*R_f_* = 0.27, 0.22, 0.15) which appeared as yellow bands, were detected among the samples obtained from inducer-treated and the wild heartwood under UV365 nm (Fig 6a). When the TLC plate was stained with 5% vanilin-H_2_SO_4_ and heated for 10 min at 100°C, all induced samples showed a blue band at *R_f_* = 0.24, a red band at *R_f_* = 0.40, and a black band at *R_f_* = 0.63. These spots corresponded to nerolidol and the wild heartwood control. Each spot presumably indicates a pure natural product or phytochemical. Each spot had a specific *R_f_* value. These results demonstrate that the resinous wood obtained by the Dodor-Wit methodwas consistent with wild heartwood’s compounds.

**Fig 5.**
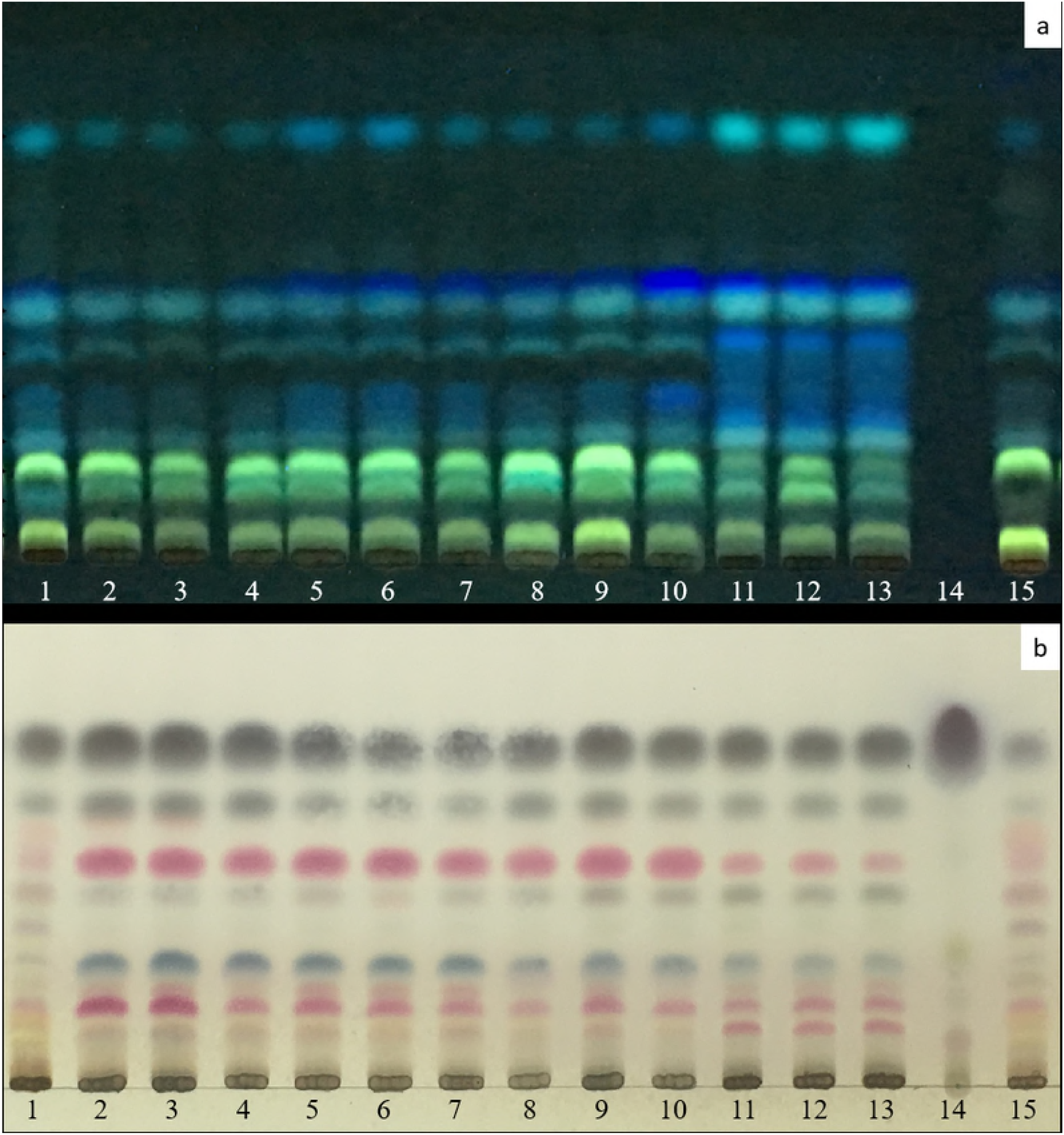
TLC chromatogram of methanol extracts of *D. odorifera* wood after application of the Dodor-Wit method. Codes 1–15 represent the 15 samples: 1 and 15, HW; 14, Nerolidol; 2–4, three replicates of LNS; 5–7, three replicates of LL; 8–10, three replicates of JF; 11–13, three replicates of BQ; (**a**) TLC chromatogram visualized under UV365 nm. (**b**) TLC chromatogram of the plate stained with 5% vanillin-H_2_SO_4_ and then heated for 10 min at 100°C.

**Fig 6.**
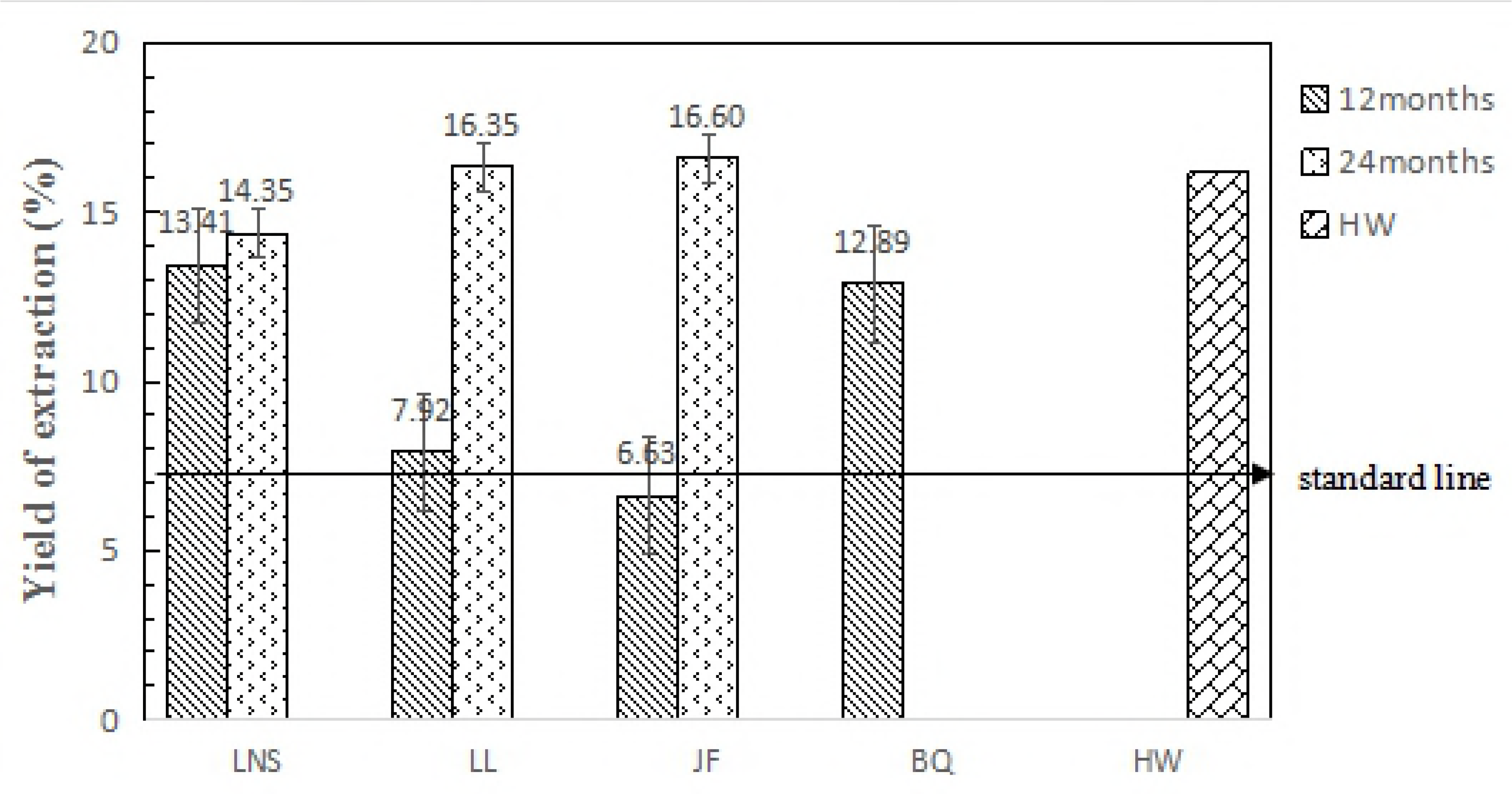
The alcohol soluble extractive content of *D. odorifera*heartwood produced by the Dodor-Wit method. Alcohol soluble extractive content was collected 12 months and 24 months after being treated in different sites. HW:natural heartwood.

### Alcohol Soluble Extractive Content

The alcohol soluble extractive content of medicinal heartwood should be no less than 8.0% according to the Chinese Pharmacopoeia [18]. The alcohol soluble extractive contents of the induced heartwood, 12 months after induction, were 13.41%, 7.92%, 6.63%, and 12.89%, for the LNS, LL, JF, and BQ plantations, respectively. The alcohol soluble extractive contents were 14.35%, 16.16%, and 16.60%, 24 months after induction, for the LNS, LL, and JF plantations, respectively. The wild heartwood was 16.16% (Fig 6).

Twenty-four months after induction, the average level of the alcohol soluble extractive content in all plantationswas 15.77% which was almost twice that required by the Chinese Pharmacopoeia, and was similar to that of natural heartwood. Twelve months after induction, the average levelof the alcohol soluble extractive content in all plantationswas 10.21%, also exceeding the Chinese Pharmacopoeiastandard. These results demonstrate that the resinous wood obtained by the Dodor-Wit24 months after inductionmethodwas consistent with natural heartwood.

### Volatile oil content and its GC fingerprint

Volatile oil content is an important index to measure the quality of heartwood of Jiang Xiang. Almost all of the oil from the induced heartwood samples was yellow and had an aroma similar to that of wild heartwood. Aside from the fact that the oil yield from sample LL-1 was 0.91%, the content of volatile oil in the induced samples (1.02–1.74%) reached the standard of the Chinese Pharmacopoeia, of ≥1% (Table 4). The content of volatile oil in the samples harvested two years after application of the Dodor-Wit method was higher than in the samples harvested after one year. The average volatile oil content per tree was 1.42%. Further analyses showed that the content of volatile oil in the natural heartwood was higher than in the induced heartwood; this is because the natural heartwood trees were older than induced trees. Generally, *D. odorifera* trees younger than 7–8 years have no heartwood and produce almost no volatile oil [20]. The yield of volatile oil in sapwood (CK) was 0.02%, which was less than that of heartwood in most of the samples (Table 3).

**Table 3.** Contents of volatile oil in the wood of *Dalbergia odorifera* trees subjected to Dodor-Wit, from different plantations in Hainan province of China. Results are presented as means ± SD.

| **sample\*** | **Contents of volatile oil (%)** | **Harvest timeafter treatment**<br>**(months)** |
| --- | --- | --- |
| **LNS-1** | 1.09±0.09 | 12 |
| **LNS-2** | 1.36±0.35 | 24 |
| **LL-1** | 0.91±0.11 | 12 |
| **LL-2** | 1.37±0.07 | 24 |
| **JF-1** | 1.45±0.18 | 12 |
| **JF-2** | 1.54±0.27 | 24 |
| **BQ-1** | 1.74±0.06 | 12 |
| **HW** | 2.89±0.79 | natural heartwood, untreated |
| **CK** | 0.02±0.02 | natural sapwood, untreated |

\* LNS, LL, JF, and BQ refer to Luoniushan plantation, Haikou; Lelai plantation, Wanning; Jianfeng plantation, Ledong; and Banqiao plantation, Dongfang. The numbers 1 and 2 after the plantation names (LNS, LF, JF, and BQ) refer to samples of wood subjected to Dodor-Wit induction, and harvested after 12 and 24 months, respectively. CK is natural sapwood. HW is natural heartwood. Each sample has three duplicates, and the result is that the content of volatile oil in three duplicates is averaged.

**Table 4.** The similarities of the nine induced-heartwood samples of *Dalbergia odorifera* to the Chinese Pharmacopoeia standard. These samples had been subjected to the Dodor-Wit method.

| JF samples* | Similarity | LL samples | Similarity | LNS samples | Similarity |
| --- | --- | --- | --- | --- | --- |
| JFA2(S2) | 93.6% | LLA2(S5) | 96.6% | LNSA2(S8) | 97.5% |
| JFB2(S3) | 97.0% | LLB2(S6) | 94.9% | LNSB2(S9) | 95.5% |
| JFC2(S4) | 97.4% | LLC2(S7) | 95.0% | LNSC2(S10) | 96.8% |
| Average | 96.0% | Average | 95.5% | Average | 96.6% |

\*The sample codes in brackets are the gas chromatography fingerprinting sample codes, which are consistent with the sample codes shown in Fig 7. The data were collected at the Luoniushan plantation (LNS), Haikou; Lelai plantation (LL), Wanning; Jianfeng plantation (JF), Ledong; and Banqiao plantation (BQ), Dongfang. A, B and C represent three replicates
of each plantation sample, and the number 2 after A, B, and C represents the harvest time of 24 months.

**Table 5.** Samples and treatments used to assess the Dodor-Wit heartwood induction method applied to *Dalbergia odorifera* trees in China.

| Sample code | Tree age<br>(years) | Sources | Harvest timeafter<br>treatment(months) |
| --- | --- | --- | --- |
| LNS-1 | 5 | Treated by Dodor-Wit | 12 |
| LNS-2 | 5 | Treated by Dodor-Wit | 24 |
| LL-1 | 5 | Treated by Dodor-Wit | 12 |
| LL-2 | 5 | Treated by Dodor-Wit | 24 |
| JF-1 | 5 | Treated by Dodor-Wit | 12 |
| JF-2 | 5 | Treated by Dodor-Wit | 24 |
| BQ-1 | 5 | Treated by Dodor-Wit | 12 |
| CK | 5 | Xylem, untreated tree<br>without heartwood | - |
| HW | ≥30 | Natural Heartwood | - |

### GC fingerprint

In order to further evaluate the quality of the artificially induced-heartwood, a volatile oil GC fingerprint of nine induced-heartwood samples (three replicates from each of three plantations) was produced, and compared with the standard chromatogram. We constructed the standard chromatogram by simulating the mean chromatogram of ten natural heartwoods. The amount of volatile oil in the inducedheartwood from the three plantations was extremely close to that of the Chinese Pharmacopoeia standard (Fig 7), with average similarities of 96.6%, 95.5%, and 96.0%, respectively (Table 4).

**Fig 7.**
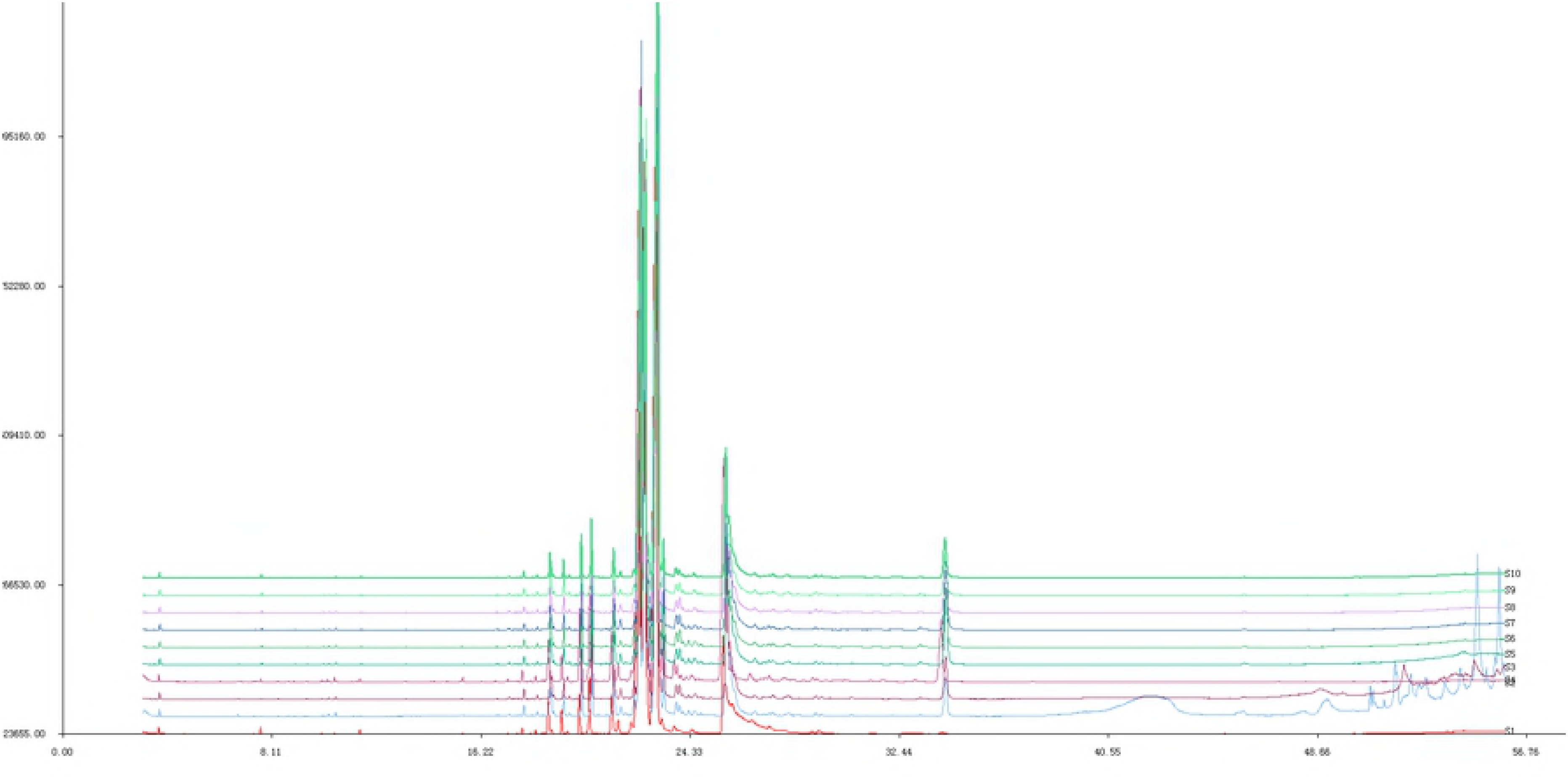
Gas chromatography fingerprints of *D. odorifera* samples treated using the Dodor-Wit method. The trees were cultivated in Hainan province of China. S1: wild heartwood; S2–S4: three batches from Jianfeng, Ledong; S5–S7: three batches from Lelai, Wanning; S8–S10: three batches from Luoniushan, Haikou.

## Materials and Methods

### Plant Material

*Dalbergia odorifera* T. Chen (Leguminosae) is indigenous in Hainan province, People’s Republic of China. The trees weresampled 12 months and 24 months after the Dodor-Wit method induction. Plant material was collected in July 2016 and July 2017 from four locations in Hainan Province: Luoniushan plantation (LNS), Haikou (110.55°N, 19.89°E); Lelai plantation (LL), Wanning (110.41°N, 18.91°E); Jianfengling plantation (JF), Ledong (108.80°N, 18.70°E); and Banqiao plantation (BQ), Dongfang (108.80°N, 18.80°E). The numbers 1 and 2 after the plantation names (LNS, LF, JF, and BQ) refer to samples of wood subjected to Dodor-wit induction, and harvested after 12 and 24 months, respectively. CK is the xylem of untreated tree without heartwood. HW is natural heartwood which was gathered from Jianfengling, Hainan Province, in July 2016. Voucher specimen: No. 20160830HNDW).

To evaluate the quality of the induced heartwood, the bark was removed from the trunk; the wood was then dried for 15 days in an oven at 40°C. The wood was then cut into small chips or flakes. The untreated CK and natural heartwood HW samples were treated in the same way.

### Dodor-Wit Treatment

The heartwood inducer is injected into the xylem of *D. odorifera* trees using a transfusion set(Fig 8). Pressure induced by transpiration causes the induction solution to infuse into the wood, causing internal wounding. The induction fluid is then transported throughout the tree within 2 to 6 h. Heartwood then forms slowly throughout the tree, without requiring follow-up treatment. The method requires sunny days with air temperatures of 30–35°C. Trees must be ≥3 years old, and the diameter at breast height (1.3 m above ground) should be ≥3 cm. The quantity of induction solution used depends on tree height, diameter of the trunk, and the size of the canopy (for sparse canopies, less solution is used).

**Fig 8.**
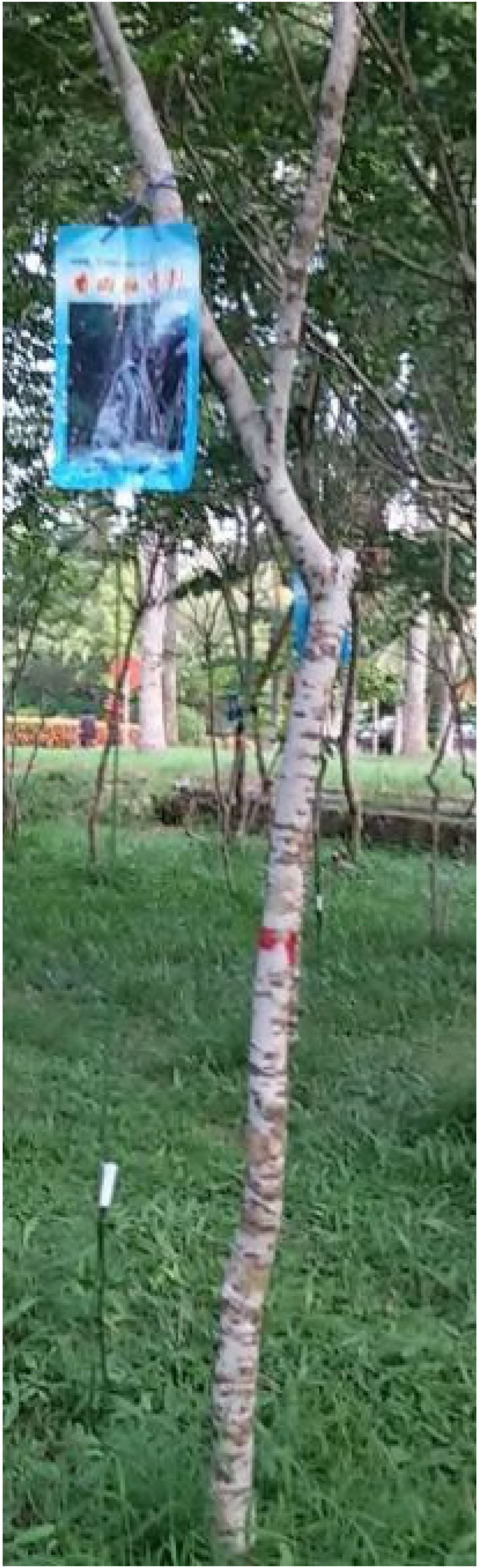
Heartwoodwas induced using Dodor-Wit methodin a 5-year-old tree of *Dalbergia odorifera*. The blue transfusion bag contains the inducer solution, which is delivered by a tube through a 5 mm hole, into the xylem capillary tissues of the tree.

The Dodor-Wit method was applied once, at the start of the experiment, and the trees were cultivated normally for the remaining period. They were felled at 30 cm above the ground. The length of the trunk (*L*) containing heartwood was measured, and the area of discoloration (*S*) was measured on the transverse plane. We measured the annual mortality rate of treated trees. The technique was tested in ten trees for each replicate group. Healthy sapwood was used as negative control (CK), and natural heartwood as positive control (HW).

In order to observe the induction process, we designed the infusion model to utilize the process of tree infusion. The dye solution comprises 5% ferric chloride and 1% acid fuchsin. Because acid fuchsin and the induction fluid migrate at the same rate, the infusion of the induction fluid can be tracked inside the trunk. Cuts are made every 30 cm above the infusion hole and the color of the cross-section is observed. From the change in color, the distance that the induction fluid has travelled up the trunk can be estimated.

### Heartwood Yield Estimation

To rapidly quantify the heartwood yield of a large number of trees treated by the Dodor-Wit method, we used a heartwood-yield estimation optimization model. The cross-section of the trunk is usually regarded as a circle, and the average width of the trunk as the diameter of a circle. Volume is then converted to the logarithm of volume. Under this approach, the induced heartwood is considered to be a cylinder and the diameter is measured at the height of 1.3 m.

To estimate the yield of heartwood produced by the Dodor-Wit method, we measured the lowest height at which induced heartwood occurred, and sawed the trunks into many 30 cm long sections, and dried them. One 30 cm section at a height of 1.3 m on the trunk was selected, and from this we cut the sapwood sample, which was then weighed. The weight of the resinous wood was then estimated according to the weight of the 30 cm stick (Fig 9).

**Fig 9.**
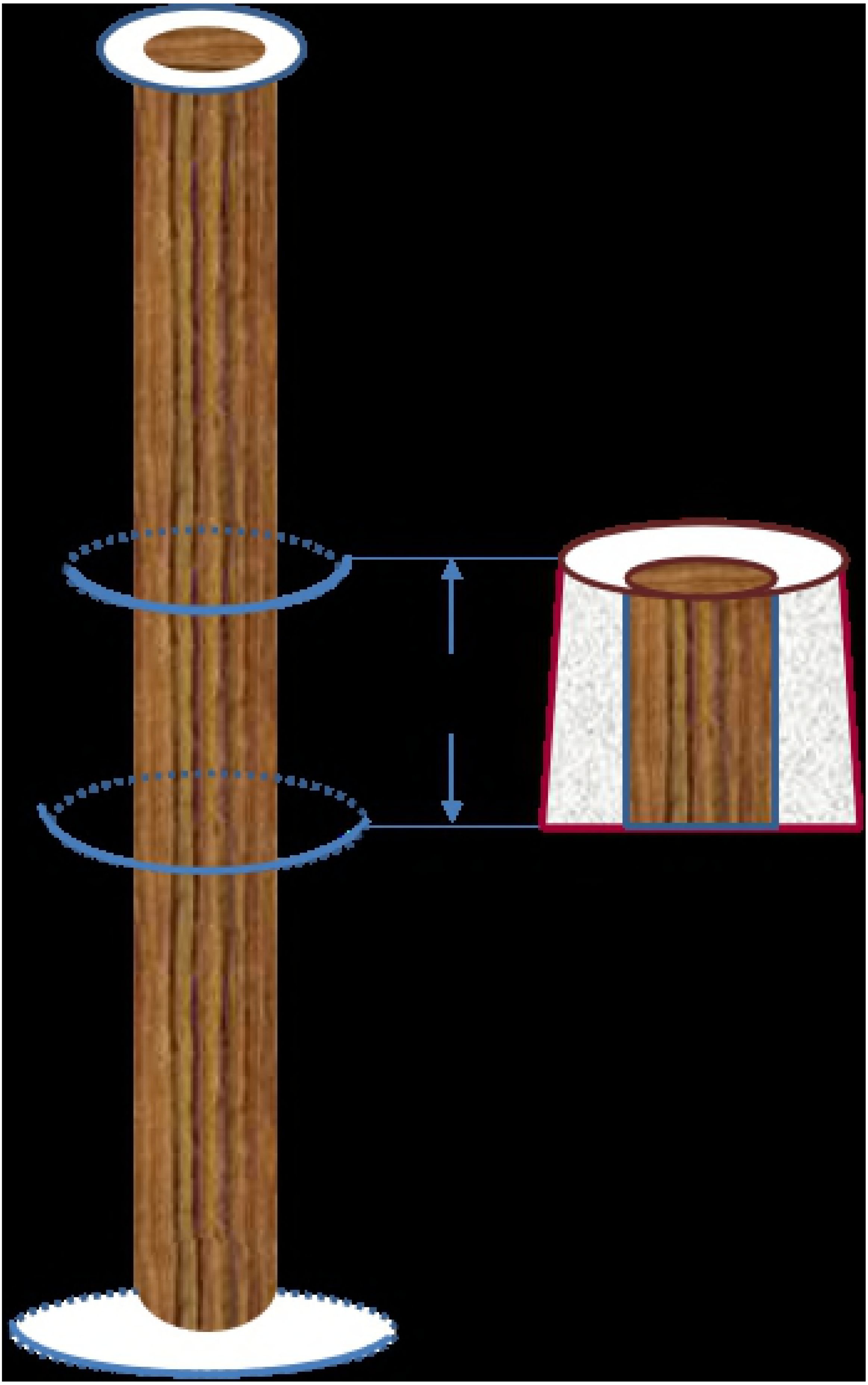
The diagram of the trunk of a *Dalbergia odorifera* tree, used to estimate heartwood yield.

The heartwood yield and heartwood ratio of each tree were calculated using the following formulae:

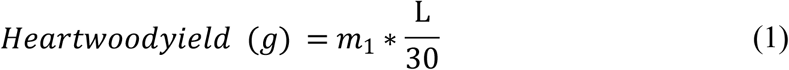

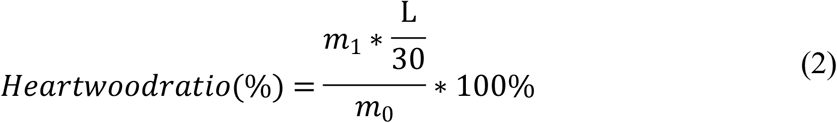

where *m_0_* represents the dry weight of all of the induced heartwood from the tree, *m_1_* represents the dry weight of the resinous wood isolated from the 30 cm stick, and *L* is the length of a trunk containing heartwood. These formulae assume that the trunk is a symmetrical cylinder, and that the ratio of resinous wood to total wood remains constant throughout the length of the trunk. For each tree, the resinous wood was separated from the white part, weighed, and the heartwood yield was calculated.

### Microscope Observations

We used a microscope(Nikon, Eclipse 80i) to study wood sections and wood powder. For the wood section method, the wood was sampled at a height of 1.3 m above ground, from natural heartwood (HW treatment), sapwood (CK treatment) and other induced trees. Small blocks of wood were then softened in a water bath at 65°C for more than 12 h. The blocks were then sliced into 15 mm thick sheets. These were placed 1.5mL centrifuge tubes containing a chloral hydrate solution (50 g of chloral hydrate, 15 mL of water, 10 mL of glycerol). This solution remains transparent for about 4 h. The sheets were observed under the microscope.

To study the wood powder, the samples were crushed and sieved using a NO. 4 sieve (250 ± 9.9 μm, 65 mesh). The sieved powder was collected. An appropriate amount of powder was placed in a 1.5-mL centrifuge tube in the same chloral hydrate solution, and observed under the microscope.

## Quality Analyses

### Thin-Layer Chromatography (TLC)

Wood samples were crushed and filtered (24 mesh sieve). Powder (1 g) was placed in methanol (10 mL) for 30 min and subjected to ultrasound (59 kHz, 500 W) using a SB25-12DTDN machine (Ningbo Xinzhi Biotechnology Co., Ltd., Ningbo, China). The solution containing the powder was then filtered, and 2 μL of the filtered solution were drawn into a capillary tube. The tube was then pressed onto a 10 × 20 cm glass TLC plate coated with silica gel (60G F_254_, Merck). The mixed solvent of methylbenzene:diethyl ether:trichloromethane (7:2:1, v/v/v) was used as the developing solvent. The TLC plate was developed and air dried. The plate was treated with 1% vanillin-sulfuric acid in EtOH, heated at 105 °C, and then viewed. Another plate containing the extracted solution was developed in solvent of methylbenzene:ethyl acetate (2:1, v/v). The TLC plates were visualized under a UV_365_ nm light.

### Alcohol Soluble Extractive Content

The alcohol soluble extractive content of the samples was detected according to the procedures given in the Chinese Pharmacopoeia[4]. Wood samples were pulverized to coarse powder (2 g), and then immersed in alcohol (100 mL) for one hour and extracted by heating reflux for one hour. After cooling and filtering, and taking precautions against loss of the solvent, the filtrate (25 mL) was evaporated to dryness in a tared flat-bottomed shallow dish, dried at 105 °C for 3 h to constant weight, and then weighed. The percentage of alcohol soluble extractive content was calculated with reference to the dried sample powder. The experiment was repeated three times.

### Volatile Oil Content

The volatile oil was extracted following the procedure given in the Chinese Pharmacopoeia [4]. The determination of volatile oil in a sample is made by distilling a sample (50 g) with water (800 mL) using specialized equipment. The distillate is collected in a tube in which the aqueous portion of the distillate is automatically separated and returned to the distilling flask. The extracted volatile oil was isolated and dried using anhydrous sodium sulfate, weighed, and stored in sealed amber flasks at –20°C until analysis. We calculated the percentage of volatile oil with reference to the dried sample powder.

### Gas chromatography fingerprinting

Volatile oils used in gas chromatography fingerprintingwere extracted by the steam distillation method. Gas chromatography was performed using an Agilent 5975C gas chromatography instrument and Agilent Chem Station software (Agilent Technologies, Palo Alto, CA, USA). Compounds were separated on a 30 m × 250 μm (length × internal diameter) capillary column coated with a 0.25μm film of 5% phenylmethylsiloxane. The injector temperature was 250 °C, helium was used as the carrier gas, and the injection volume (n-hexane solution) was 1 μL. The temperature program started at 50 °C for 2 min, increased by 6 °C min^−1^ to 135°C, and was held there for 1 min, then increased at 1 °C 20 min^−1^ to 152 °C, and was held there for 2 min. The temperature was then increased at 2 °C min^−1^ to 172 °C, then at 10 °C min^−1^ to 250 °C, and was held there for 2 min. The “*Similarity evaluation system for chromatographic fingerprint of TCM* (Version 2004 A)” was used to analyze experimental data in the experiment.

## Conclusions

The Dodor-Wit method relies on pressure induced by transpiration in *D. odorifera* trees to infuse the induction fluid into the trunks, roots, and lateral branches, andthen induce heartwood prodution throughout the tree. The results show the heartwood can be induced in *D. odorifera* trees of 5–8 years old,rather than the decades required for it to form naturally. 1–2 years after treated by the Dodor-Wit method, the induced heartwood quality can reach the standard of Chinese Pharmacopoeia. Although there are some methods for inducing heartwood in *D. odorifera*, such as drought, microorganism, phytohormone and so on, but they are local induction, the yield of induction is low, the experimental samples are small. We have been doing this research for more than ten years in several plantations in Hainan, Guangdong, Guangxi. The inductive technical system includes tree selection, environmental conditions, operating procedures, etc. Through a large number of sample data, induction technique can effectively induce heartwood production in cultivated *D. odorifera*, which yield and quality stability, and easy to operate. So it is a commercially feasible technique to produce medicinal heartwood. The combination of this method with large-scale cultivation of *D. odorifera* trees could also meet the demand for heartwood and its essential oils. The application of Dodor-Wit in cultivated trees of *D. odorifera* to produce valuable heartwood may help indirectly to conserve wild *D. odorifera* resources.

## Acknowledgments

This work was supported by the CAMS Initiative for Innovative Medicine (CAMS-I2M) (2016-I2M-2-003), the key Research Project of Hainan Province, China (Grant No. ZDYF2018123), and the Program for Creative Research Groups of Hainan Provincial Natural Science Foundation (No. 2017CXTD022).

**Author Contributions**
Hui Meng was a major experimenter and author of paper; JianHe Wei designed experiments, manuscript revision andfunding acquisition; and Yun Yang participated in the experimental design, all experiments and writing-original draft preparation; DeLi Chen and Zheng Zhang participated in experimental design, implementation and helped analysis and editing; Xiangsheng Zhao established the method of GC fingerprint. Yangyang Liu, Jian Feng and Peiwei Liu helped analyze the results.

## References

1. Dezhao C, Bangyu Y, Yunyi F, Chaozong Z, Ruohui Z, Chensen D, et al. Leguminosae. Flora of China, Volume 40. Zhi W, editor. Beijing: Science Press; 1994. p. 114.in Chinese.

2. Delin W, Zhaofen W, Bangyu C, Yunfei D. Papilionaceae. Flora of Guangdong, Volume5. Telin W, editor. Guangdong: Guangdong Science and Technology Press; 2003. p. 225.in Chinese.

3. GB/T 18107-2017. Hongmu. Beijing: China Standard Press; 2017.

4. Pharmacopoeia Commission of the People’s Republic of China. Pharmacopoeia of the People’s Republic of China, Part 1. Beijing: China Medical Science Press; 2015. p. 229–230.

5. Tentative standard of new drug of Chinese medicine: YBZ04332003-2008Z

6. Commission of Chinese Pharmacopoeia. Chinese Pharmacopoeia. Fragrant Ingredients Pain-Re, Part 1. Beijing (China): Chinese Medicine Science and Technology Publishing House Press; 2015. p. 447–448.

7. Pharmacopoeia Commission of the People’s Republic of China. Pharmacopoeia of the People’s Republic of China: Guanxin-Dan-shen pills. China Medical Science Press: Beijing, China, 2015; Part 1, p. 1310.

8. IUCN. IUCN Red List of Threatened Species. IUCN Global Species Programme Red List Unit: Cambridge, 2008. Available from: http://www.iucnredlist.org (cited 3 April 2009).

9. Lin L, Qiu JY, Cai YW, Cai C, Li XM, Wang KY, et al. Production status, problems and countermeasures of rare southern medicinal species in Guangdong. Chinese Modern Traditional Medicine. 2013;15:127–130.in Chinese.

10. Wu YB, Zai CH, Zhuang XY, Liang SY, Wu BL. Early growth of the plantations of *Dalbergia odorifera* in Zhaoqing, Guangdong. Guangdong Forestry Science and Technology. 2010;26:36–40.in Chinese.

11. Guo WF, Jia HY. The introduction of *Dalbergia odorifera* in southern subtropical area of Guangxi. Journal of Fujian Forestry Science and Technology. 2006;4:152–5.in Chinese.

12. Xu GY, Duan ZL. Research on the introduction of technology of *Dalbergia odorifera* at Menghai County. Ningxia Journal of Agriculture and Forestry Science and Technology. 2014;55:24–5.in Chinese.

13. Yao QH, Lin QJ, Zheng ZL. Study on *Dalbergia odorifera* suitable introduction regions in Fujian Province. Journal of Fujian Forestry Science and Technology. 2012;39:89–94.in Chinese.

14. Zou XH, Zhang JL, Qiu XH, Xiao YH, Liu LS. Report of introduced cultivation and afforestation experiment of *Dalbergia odorifera*. Jiangxi Forestry Science and Technology. 2012;3:37–8.in Chinese.

15. Chen QX, Li XW, Wang JW, Li KE, Xia HT. Introduction of *Dalbergia odorifera* T. Chen in Wenzhou and trial of afforestation. Forest Sci Technol. 2015;3:34–5.in Chinese.

16. Huang JH. A Preliminary report on afforestation with *Dalbergia odorifera* in Yibin. Journal of Sichuan Forestry Science and Technology. 2018;39:37–9.in Chinese.

17. Meng H, Yang Y, Feng JD. Status and development of introduction and cultivation of *Dalbergia odorifera*. Guangdong Agricultural Sciences. 2010;7:79–80.in Chinese.

18. Taylor AM, Gartner BL, Morrell JJ. Heartwood formation and natural durability - a review. Wood Fiber Sci. 2002;34(4), 587–611.

19. Chan JM, Raymond CA, Walker JCF. Development of heartwood in response to water stress for radiata pine in Southern New South Wales, Australia. Trees Struct Funct. 2013;27:607–617. doi: 10.1007/s00468-012-0815-3.

20. Jia RF. Study on the artificial promotion of heartwood formation by flavonoids. Doctoral thesis, Research Institute of Tropical Forestry, Chinese Academy of Forestry, Guangzhou, People’s Republic of China, 7 July 2010.

21. Wilkins AP. Sapwood, heartwood and bark thickness of silviculturally treated *Eucalyptus grandis*. Wood Sci Tech. 1991;25:415–423.

22. Wei JH, Meng H, Yang Y, Feng JD, He MJ, Zhang Z. A method of reducing Jiang Xiang formation in *Dalbergia odorifera* T. Chen. China patent ZL200910192294. 2010.

23. Lin L, Wei M, Xiao S, Xu X, Hu Z, Qiu J, et al. The influence of external stimulation on content and quality of volatile oil in Lignun Santali Albi. Journal of Chinese Medicinal Materials. 2000;3:152–4.in Chinese.

24. Lin QY, Cai YW, Yuan Liang, Zhong YQ, Qiu JY, Lu AN, et al. Experimental study on external stimulation of Lignum Santali Albi. Journal of Chinese Medicinal Materials. 2000;23:437–8.in Chinese.

25. Meng H, Yang Y, Chen B, Chen WP, Feng JD, Wei JH. External stimulation of the formation of Heartwood in *Dalbergia odorifera*. The Eighth National Symposium on Medicinal Plants and Botanical Drugs, Hohhot, Inner Mongolia, People’s Republic of China, 2009;86.

26. Liu XJ, Xu DP, Yang ZJ, Zhang NN. Effects of plant growth regulators on growth, heartwood formation and oil composition of young *Santalum album*. Scientia Silvae Sinicae. 2013;49:143–9.in Chinese.

27. Zhou SQ, Huang X, Zhou Y. A biological method for inducing red sandalwood odorization. China patent, CN201410104193.5, 2014.

28. Sun SS, Meng ZX, Zhang DW, Guo SX. Field screening for fungus inducing active substances from *Dalbergia odorifera*. Chin Pharm J. 2017;52:227–283.

29. Chen XY, Liu YY, Yang Y, Feng J, Liu PW, Sui C, et al. Trunk surface agarwood-inducing technique with *Rigidoporus vinctus*: an efficient novel method for agarwood production. PLoS ONE. 2018;13:e0198111. DOI: 10.1371/journal.pone.0198111.

30. Liu YY, Chen HQ, Yang Y, Zhang Z, Wei JH, Meng H, et al. Whole-tree agarwood-inducing technique: an efficient novel technique for producing high-quality agarwood in cultivated *Aquilaria sinensis* trees. Molecules. 2013;18:3086–3106. doi: 10.3390/molecules18033086.

